# Krüppel-Like Factor 14 Improves Cardiac Function After Myocardial Infarction

**DOI:** 10.1101/2020.03.25.008052

**Authors:** Yating Wang, Danyan Xu, Li Shen

## Abstract

**Background:** Reversal of heart failure after myocardial infarction (MI) remains a global challenge. The goal of a clinical physician is to prevent heart failure immediately following the infarct. Since traditional strategies are only partially helpful for improving cardiac function, there is an urgency to identify novel approaches to maintain cardiac function.

**Objectives:** The aim of this study was to determine the role of Krüppel-Like Factor 14 (KLF14) in the heart post-MI.

**Methods and Results:** Wild-type and KLF14 overexpression mice were subjected to MI. At four weeks following MI, the infarct size was significantly reduced, left ventricular fraction shortening was significantly preserved, and left ventricular chamber dilation was attenuated in the KLF14 overexpression group. Transmission electron microscopy studies revealed striking evidence of decreased mitochondrial damage in the KLF14 overexpression myocardium following MI. To gain insight into the underlying relationship between KLF14 and mitochondria, we examined a broad array of genes involved in mitochondrial function and found that KLF14 overexpression was involved in the decreased apoptosis of mitochondria induced by H_2_O_2_. This beneficial effect was associated with increased levels of PGC-1α.

**Conclusions:** These results demonstrate that KLF14 plays a critical protective role in the heart after MI by regulating mitochondrial function, which may be achieved by regulating PGC-1α.

## Introduction

As the leading cause of left ventricular dysfunction, heart failure is a major cause of morbidity and mortality throughout the world [1, 2]. Although advanced clinical and pharmacological interventions are currently available, the post-discharge mortality and rehospitalization rates within 69 − 90 days in patients with heart failure remain high, which can reach up to 15% and 30%, respectively [3]. To maintain contractile function, ATP supply is the most important for the heart. As the crucial organelle for ATP, mitochondria generate 95% of cardiomyocyte ATP from the oxidative metabolism of fatty acids and carbohydrates [4]. Small changes in mitochondrial function can have huge impact on heart function because it is critical to maintain cardiac function by a continuous ATP supply, which is produced within the mitochondria [5]. Increasing evidence has shown that during myocardial ischemia, mitochondrial homeostasis, including FAO, OXPHOS, mitochondrial biogenesis, dynamics, and autophagy are disrupted, resulting in reduced energy production, decreased myocardial contractility, increased cell death, and the development of heart failure during the process [6,7]. Therefore, exploring the molecular mechanisms that regulate mitochondrial health may help to treat heart failure in the future.

Kruppel-like factors (KLFs) represent a family of zinc finger transcription factors, which include 17 siblings (KLF1-17). Acting as transcriptional repressors or activators, KLFs are expressed among different tissues. KLF protein disorders are involved in the pathogenesis of cardiovascular disease, such as cardiac development, cardiac hypertrophy, and atherosclerosis. Recently, researchers have shown that KLFs play a critical role in regulating the number and function of mitochondria. Mallipattu et al. reported that KLF6 binds to SCO2, a promoter of one key mitochondrial gene. In addition, the mice develop kidney failure in response to renal toxins by reducing the mitochondrial complex size and density when KLF6 is lacking in kidney podocytes [8]. In addition, KLF11 has been identified as a new regulator of mitochondrial function in adipose tissue, and is required for the “browning” of white fat cells into mitochondria-enriched and energy-consuming cells [9]. In addition, Liao et al. suggested that KLF4 is required for mitochondrial biogenesis and homeostasis in the heart [10]. In cultured pulmonary artery smooth muscle cells, KLF5 drives survivin expression, generates mitochondrial hyperpolarization, and protects against apoptotic cell death [11]. In *Caenorhabditis elegans*, KLF3 is critical for fatty acid beta-oxidation related enzyme expression in the mitochondria [12]. This suggests that KLFs play a crucial role in the regulation of mitochondria function.

KLF14 is a member of the KLF family located in the human chromosome 7q32.2 and encodes 323 amino acids, [13]. KLF14 is highly expressed in the heart [14]. Genome-wide association studies (GWAS) have shown that single nucleotide polymorphisms (SNPs) in the Klf14 locus decreased HDL-C and increased TG in European populations [15]. Subsequently, in vivo studies using apoe^−/−^ mice showed that the lack of KLF14-mediated activation of the apoA-I promoter in the lever reduced the levels of HDL-C and cholesterol efflux, which significantly enhanced the development of atherosclerotic lesions. Recent GWAS has confirmed that a Klf14 variant (rs4731702) is associated with the etiology of atherosclerotic cardiovascular disease (ASCVD), which includes ischemic stroke and myocardial infarction (MI) [16]. In addition, the authors suggest that Klf14-rs4731702 and missense Klf14-rs111400400 may represent two different causal risk factors. Thus, the Klf14 gene both regulates metabolism, and is also associated with atherosclerotic-related diseases, such as MI. These findings suggest that KLF14 may have pleiotropic effects on non-lipid phenotypes that pose a risk of ASCVD. However, the role and function of KLF14 in MI remain poorly defined. Klf14 is a maternally-imprinted gene expressed from the maternal allele in both embryonic and extra-embryonic tissues [17, 18]. In addition, it has been established that mitochondria are matrilineally inherited organelles. Therefore, whether KLF14 participates in mitochondrial function in the heart and the associated mechanisms are important to elucidate.

## Methods

### Animals

The animal protocols were reviewed and approved by the Ethical Committee of the Second Xiangya Hospital of Central South University. A cohort of C57BL/6 male mice (6 weeks old) was selected for the study and housed on a 12-h light/12-hdark cycle in a temperature-controlled environment (22°C) with free access to standard laboratory chow and tap water.

### Myocardial infarction

As previously described, left anterior descending (LAD) ligation is the most common model used to induce myocardial infarction [19]. This model was selected because myocardial infarction is the underlying cause in nearly 70% of patients with heart failure. The mice were anaesthetized with 1% phenobarbital, endotracheally intubated, and mechanically ventilated with room air (respiratory rate of 100 breaths/min and tidal volume of 250 μL). A left thoracotomy was performed between the third and fourth ribs and the LAD was ligated. The animals were monitored daily after surgery. A left thoracotomy with equal procedure duration to that of the heart failure group, but without LAD ligation, was undertaken in the sham group (control).

### Echocardiography

The mice were anesthetized with phenobarbital and monitored by transthoracic two-dimensional echocardiography with a 12-mHz probe. By performing an M-mode interrogation in the parasternal short-axis view at the level of the greatest LV end-diastolic dimension, LV end-diastolic dimension (LVEDD) and LV-end-systolic dimension (LVESD) were acquired. These data can be used to calculate the percentage of fractional shortening (FS), which was calculated using the following formula: [(LVEDD-LVESD)/LVEDD] *100.

### Determination of the LV infarct size

The myocardial infarct size was determined as previously described [20]. Specifically, the mice were euthanized and the heart was quickly harvested and sliced into five 1.0 mm-thick sections perpendicular to the long axis of the heart. Then, 1% triphenyltetrazolium chloride (Sigma) was used to incubate these sections for 30 min at room temperature. The incubated sections were subsequently photographed.

### Isolation of mouse heart mitochondria

Mouse heart mitochondria were isolated as previously described [21]. Briefly, the mice were anesthetized by phenobarbital, and the heart was quickly excised. The heart was minced on ice, resuspended in 1 mL of homogenization buffer (10 mmol/L tris, 250 mmol/L sucrose, 1’ protease inhibitor, pH 7.5) supplemented with 1 mmol/L EDTA, and homogenized with a glass Dounce homogenizer and Teflon pestle. The homogenates were centrifuged at 1000’ g for 5 min at 4°C. The supernatants were recentrifuged at 10 000’ g for 15 min to pellet the mitochondria. The pellet was washed twice with homogenization buffer.

### Klf14 overexpression

A flag-tagged (N-terminal) open reading frame of human Klf14 was cloned into the vector (under the control of a TNT-promotor) and was subsequently used for Adeno-Associated Virus 9 (AAV9) generation. Adenovirus generation was performed as previously described [22]. An intravenous injection of AAV9-hKLF14-flag-GFP (1 × 10^13^vg/mL, 100 μL) was performed in 6-week-old male mice by tail vein injection four weeks before the onset of permanent MI. In vitro adenovirus (1 × 10^9^ pfu) infection was performed on day 7 after cardiomyocyte isolation.

### Cell culture and treatment

The hearts were digested in Trypsin (2−3 mg/mL, Sigma) by serial digestion and pre-plated twice to remove fibroblasts. Cardiomyocytes were seeded at a density of 1000 cells/mm^2^ on plastic culture plates (BD Falcon) and kept at 37°C in Dulbecco’s modified Eagle’s medium (DMEM, Gibco) supplemented with 10% (v/v) heat-inactivated fetal bovine serum (FBS), penicillin (P, 100 U/mL, Gibco), streptomycin (S, 100 µg/mL, Gibco), and 5-bromo-2-deoxyuridine (BrdU, 100 μmol/L, Sigma). Prior to treatment, the medium was changed to serum-free DMEM for 16 h. At 72 h following adenovirus infection, the cells were subjected to H_2_O_2_ treatment (200 μmol/L) for 24 h [23].

### Histological analysis and transmission electron microscopy (TEM) analysis

The mouse hearts were fixed in 4% paraformaldehyde for 24 h, dehydrated in increasing concentrations of ethanol, and embedded in paraffin. Heart sections (5 μm) were stained using Masson trichrome (Sigma-Aldrich). A Nikon Eclipse 80i microscope and NIS Elements software were used to capture the images. Transmission electron microscopy was used to obtain a closer observation of hepatocyte mitochondria. The heart tissues were cut into 1-mm^3^ sections on ice and fixed in a 2% glutaraldehyde solution. The samples were re-fixed using osmic acid for a few hours, observed, and photographed via a transmission electron microscope.

### RNA isolation and quantitative real-time PCR

Total RNA from the heart and cardiomyocytes was isolated using Trizol reagent (Invitrogen) according to the manufacturers’ instructions, and was quantified using a NanoDrop Spectrophotometer (Rockland, DE, USA). Complementary DNA (cDNA) was synthesized using a cDNA Synthesis Kit (Thermo, Waltham, MA, USA). RT-PCR was performed with SYBR Green qPCR Master Mix reagent (Bimake, Houston, TX, USA) in a 7300 Real-Time PCR System (Applied Biosystems), according to the manufacturers’ instructions. For each gene, RT-PCR was run on five samples in each group, with each sample in triplicate. All samples were quantified using comparative CT methods for the relative quantification of gene expression, normalized to 18SrRNA. The quantitative real-time PCR primer sets are shown in Supplemental Table 1.

### SDS-polyacrylamide gel electrophoresis and immunoblotting

Western blots were performed according to the standard technique. Briefly, homogenates of the heart tissue, isolated mitochondria, and cultured cardiomyocytes were subjected to 12% SDS-PAGE and transferred to polyvinylidene fluoride (PVDF) membranes (Millipore). After blocking in 5% defatted milk for 2 h, the membrane was probed with a primary antibody overnight at 4°C. The blots were subsequently incubated with horseradish peroxidase-conjugated secondary antibodies. Chemiluminescence was detected using an Amersham Imager 600. The total level of protein was normalized to that of Cox IV or tubulin.

### Statistical analysis

The results of the data analysis are presented as the mean ± SEM. The statistical comparison between the two groups was performed using unpaired *t-*tests. Statistical analyses were performed using GraphPad Prism software version 6. A *p* value < 0.05 was considered to be statistically significant.

## Results

### KLF14 in the mice hearts post-MI

Using a mouse model of permanent coronary occlusion, changes in KLF14 expression after evaluating MI. KLF14 mRNA expression was low in the infarcted tissue in the heart four weeks post-MI (Fig. 1A). Consistent with this result, a Western blot confirmed that KLF14 expression was decreased in the heart tissue of the infarction area (Fig 1B). Given the importance of mitochondria in cardiac function, it is reasonable to assume that KLF14 is an altered mitochondrial protein. We found that there was a robust decrease in mitochondrial KLF14 expression in the infarcted tissue (Fig. 1C). These findings suggest that KLF14 plays an important role in the heart post-MI.

**Figure 1.**
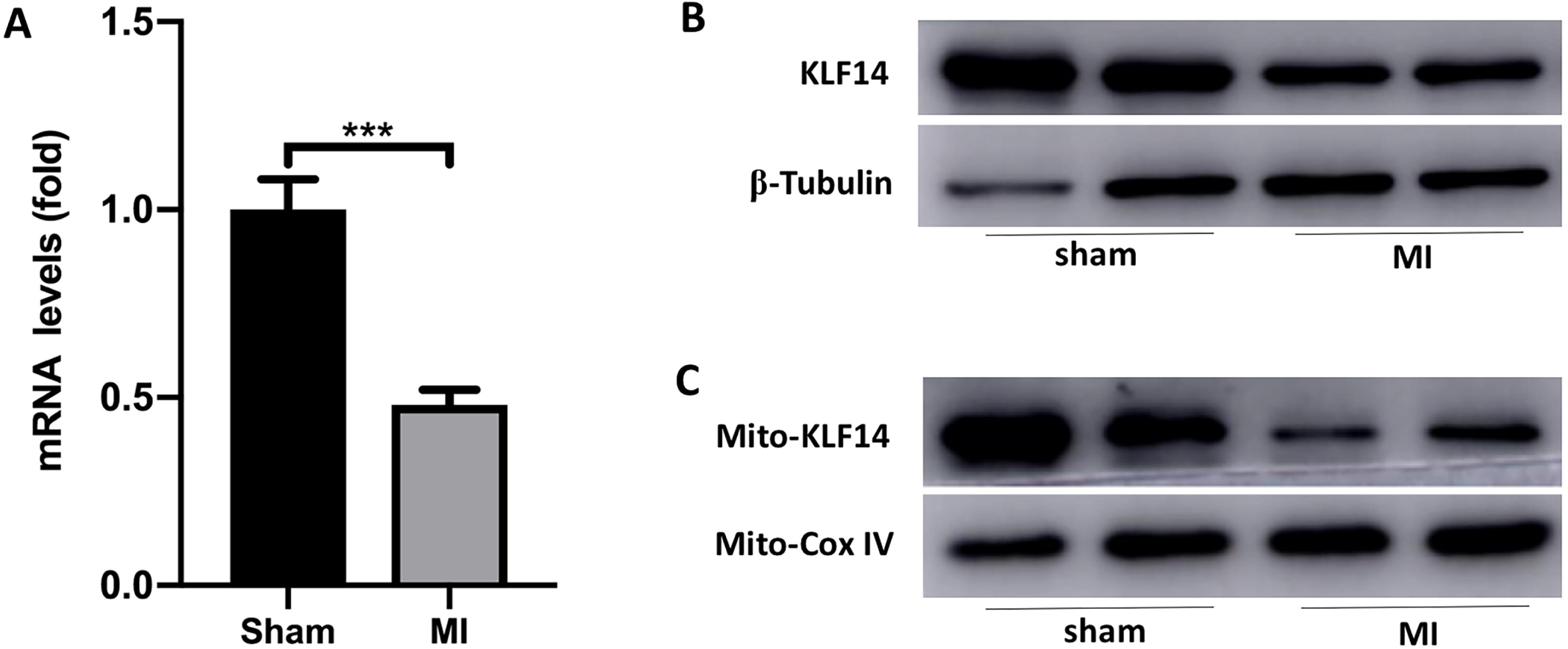
KLF14 decreased in the mice heart tissue post-MI. (A) KLF14 mRNA levels and (B) protein expression of sham and four-week post-MI LV tissue. (C) Mitochondrial KLF14 protein expression was analyzed by immunoblotting of mitochondria protein lysates from the heart tissue of sham and four-week post-MI mice. ^***^*P* < 0.001.

### AAV9 overexpression of KLF14 improves cardiac function after MI in mice

To address the role of KLF14 in MI, we determined the effects of AAV9-mediated overexpression of KLF14 on cardiac function after MI in mice. At six weeks, the mice were subjected to an intravenous tail vein injection. To assess whether KLF14 had been induced, we assessed KLF14 mRNA levels in mice. In the overexpression group, the level of KLF14 mRNA in the heart was significantly increased (Fig. 2A). There was also flag protein expression (Fig. 2B) four weeks after the intravenous tail vein injection. To determine the effect of KLF14 overexpression on the infarct size, the mice were subjected to MI at 10 weeks of age (four weeks after the intravenous tail vein injection). The infarct size was determined on day 2 after MI with the use of triphenyl tetrazolium chloride-stained heart sections. We found that the infarct size was significantly decreased in the KLF14 overexpression group in comparison with the control group (Fig. 2C). To determine the role of KLF14 on cardiac function post-MI, transthoracic two-dimensional echocardiography was performed after four weeks of MI. The KLF14 overexpression mice had a lower increase in the end-diastolic and end-systolic dimensions in comparison to the control group. This was associated with decreased LV dysfunction, as reflected by a less reduced LV fractional shortening (Fig. 2D). The hearts were excised at four weeks after MI and Masson trichrome staining was performed. We found lower fibrosis and scar expansion in the KLF14 overexpression mice compared to the control group (Fig. 2E). Furthermore, the overexpression of KLF14 in the mice markedly inhibited the heart failure-induced activity of B-type natriuretic peptide (Fig. 2F). These results revealed that KLF14 overexpression contributed to helping the heart survive from heart failure post-MI.

**Figure 2.**
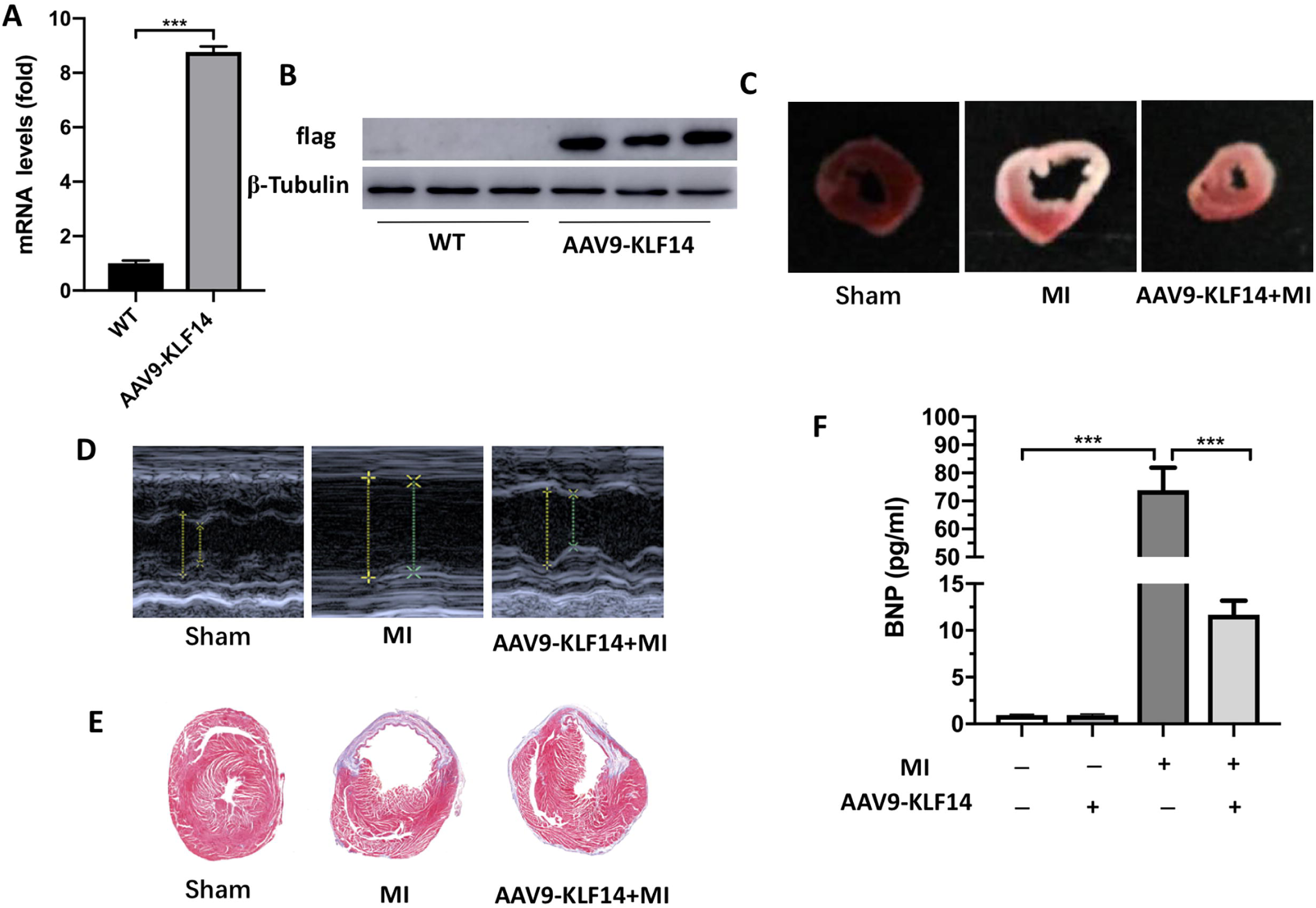
KLF14 overexpression improves cardiac function post-MI. Four-week-old mice were injected with either AAV9-hKLF14-flag-GFP or AAV9-control-GFP via the tail vein. Four weeks after injection, qPCR and a Western blot were performed on heart tissue for the levels of hKLF14 mRNA (A) and flag protein expression (B) separately. The cardiac infarct size of each group was measured by TTC (C). Left ventricular function was assessed by transthoracic echocardiography, and FS values were determined for cardiac function (D). Representative images of LV remote myocardium in WT and KLF14 overexpression mice stained with Masson trichrome four weeks post-MI (E). Plasma BNP levels (F) were measured in WT and KFL14 overexpression mice at four weeks post-MI. ^***^*P* < 0.001.

### KLF14 overexpression improves mitochondrial function

As previously described, we observed a significant reduction of KLF14 in the myocardial mitochondrial protein post-MI, which indicated that there may be a correlation between KLF14 and mitochondria. Transmission electron microscopy studies revealed striking evidence of decreased mitochondrial damage in the KLF14 overexpression myocardium following MI, including mitochondrial disarray, degeneration, fragmentation, elongation, and increased heterogeneity (Fig. 3A). To gain insight into the underlying relationship between KLF14 and the mitochondria, we expressed Ad-hKlf14 in cardiomyocytes in vitro, and confirmed the expression by Western blot (Fig. 3B). To explore how mitochondrial KLF14 overexpression in cardiomyocytes respond to hypoxia, we selected exogenous (hydrogen peroxide [H2O2]) oxidative stress and then examined a broad array of genes involved in mitochondrial function, including transcription regulation of FAO, OXPHOS, dynamics, apoptosis and autophagy. In addition, we found that there were significant changes in the apoptosis genes (Fig. 3C). To verify the role of KLF14 participating in apoptosis, expression of the pro-apoptotic Bax and the antiapoptotic Bcl-2 were measured. H2O2 increased Bax while decreasing Bcl-2 expression, and these H2O2-evoked changes were inhibited in KLF14-overexpressing cardiomyocytes (Fig. 3D). These findings indicate that KLF14 promotes cardiac function post-MI via regulating mitochondrial function, especially the apoptosis of mitochondria.

**Figure 3.**
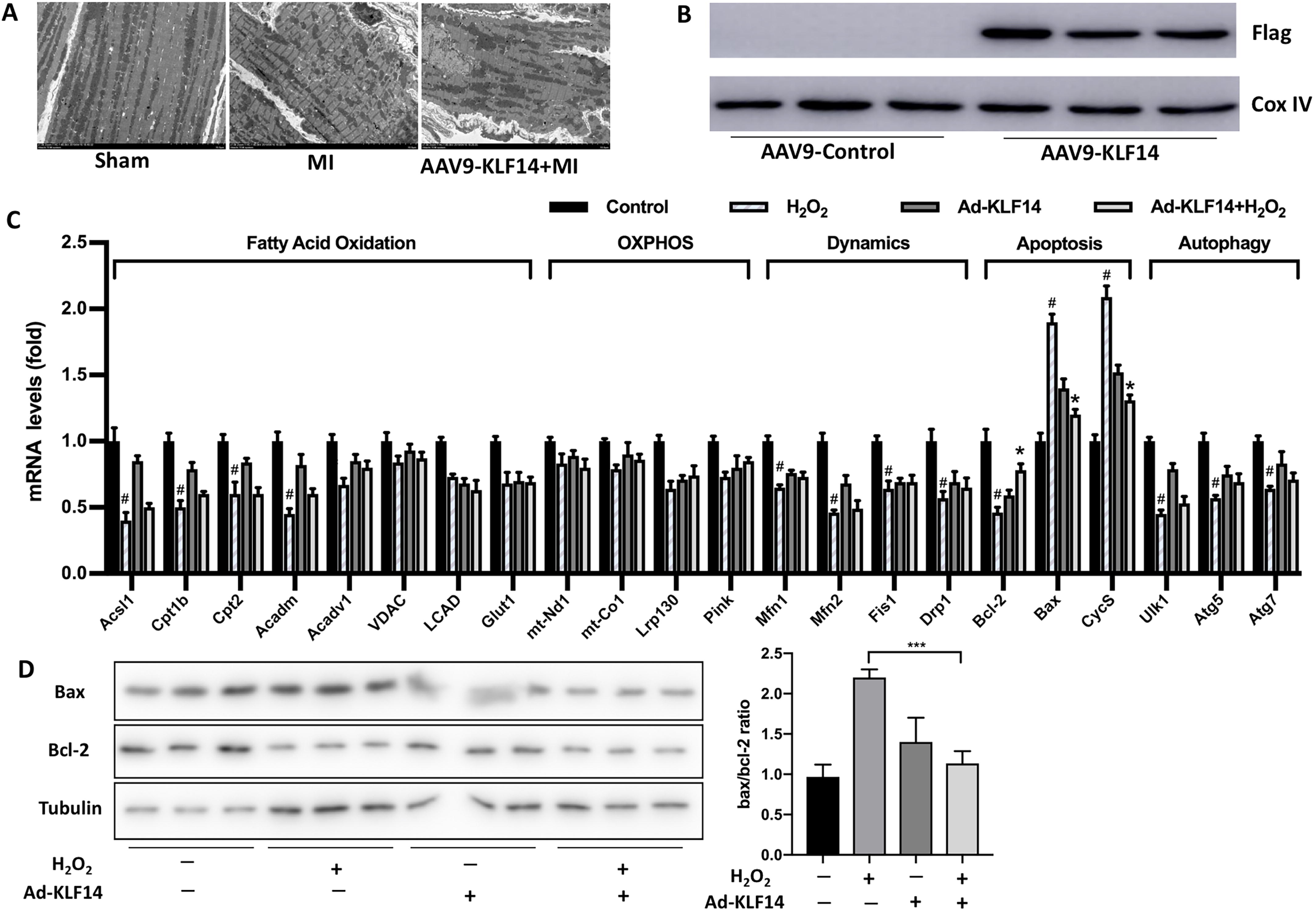
KLF14 overexpression improves mitochondrial function. The myocardium ultrastructure, as assessed by transmission electron microscopy (TEM), showing mitochondrial damage after MI (A). Detection of hKFL14 protein by performing a Western blot on cardiomyocytes for flag protein expression (B). Treatment of cultured cardiomyocytes with H_2_O_2_ in the absence or presence of hKFL14 overexpression, expression of some metabolic and mitochondrial genes was measured by qPCR (C), and Bax and Bcl-2 expression was detected by Western blot (D). #*P* < 0.05 between the control and H_2_O_2_ groups. ^*^*P* < 0.05 between H_2_O_2_ and Ad-KLF14+H_2_O_2_ group.

### KLF14 overexpression improves the levels of PGC-1α mRNA

Numerous factors can affect mitochondrial function. To gain insight into the underlying molecular mechanisms, we assessed a broad spectrum of genes that are involved as coactivators of mitochondrial biogenesis, nuclear respiratory factors, mitochondrial transcription factors, and mtDNA replication factors. We also found that the level of PGC-1α mRNA was significantly increased in the KLF14-overexpressed heart (Fig. 4), suggesting the possibility that KLF14 may regulate mitochondrial genome transcription.

## Discussion

This study is the first to demonstrate that KLF14 overexpression improves cardiac function and decreases heart remodeling after MI in response to a permanent left anterior descending artery occlusion, which may be achieved by regulating PGC-1α.

Heart failure typically occurs after MI since the heart undergoes a remodeling process after myocardial infarction, which includes LV dilatation and systolic dysfunction. In many patients, especially those with large infarcts, strategies for preventing remodeling are usually centered on a neurohormonal blockade and often fail to prevent heart failure. Herein, we identify KLF14 as a novel and potent inhibitor of heart failure post-MI. KLF14 overexpression leads to substantially decreased dilatative remodeling and better-preserved LV function.

Over time, the loss of myocardial cells, even if the incidence is very low, can lead to the collective death of myocardial cells, and has been reported to cause cardiac dysfunction and heart failure following myocardial infarction [24]. Moreover, Bcl-2 is a known anti-apoptotic factor. Bcl-2 overexpression can protect the body from oxidative stress-induced apoptosis and retain cardiomyocyte viability and LV function during ischemic [23, 25]. Bcl-2, especially the Bcl-2/Bax ratio, is an upstream regulator of mitochondrial cytochrome c (CycS) release and is an important determinant of the susceptibility of the cells to apoptosis. Our findings suggest that KLF14 overexpression leads to increased Bcl-2/Bax, which may be the mechanism by which cardiac function is improved after MI. The lower accumulation of the fragmented/damaged mitochondria in the KLF14-overexpressing heart may suggest that KLF14 also affects the process of apoptosis.

It has been suggested that a reduction of PGC-1α is associated with the development of heart failure. To date, the transcriptional regulation of mitochondrial function in the heart is largely due to the PGC-1α transcriptional module, which plays an integrative role in relaying developmental and physiological signals to a key subset of transcription factors to guide the coordinated expression of nuclear and mitochondrial genes [26]. We believe that our data demonstrates that KLF14 is a novel and essential component of this transcriptional module, and the basis for this conclusion is obvious. KLF14 overexpression showed a stark contrast to the mitochondrial phenotype observed in the Ppargc1a KO animals, including mitochondrial and cardiac function.

However, there are limitations associated with this study, and there are several aspects of KLF14 in cardiac and mitochondrial biology that remain incompletely understood and will require further investigation. First, these findings are limited to the overexpression of KLF14 by AAV9. These results will be more convincing if Klf14-KO mice are studied. Second, the method by which KLF14 interacts with PGC-1α is not completely understood. Third, whether a dose of KLF14 regulates a unique subset of mitochondrial genes remains to be clarified.

In summary, our current work provides cogent evidence implicating KLF14 as a negative transcription factor that is indispensable for the heart’s response to ischemia. Collectively, these observations contribute to the increasing appreciation that KLFs are the key molecular regulators of cardiac biology.

## Abbreviation

MI: myocardial infarction
HDL-C: high-density lipoprotein cholesterol
TG: triglyceride
GWAS: Genome-wide association studies
ASCVD: atherosclerotic cardiovascular disease

## Acknowledgments

The authors express our appreciation to Minzhi Ouyang, Zefang Peng, Yun Zhu and the Echocardiography Core for their expert assistance.

## Authors’ contributions

Yating Wang and Li Shen designed and conducted the study and analyzed the data. All authors contributed intellectually to data interpretation and manuscript preparation.

---

**Figure 4.** KLF14-induced improvement of mitochondrial function in cardiomyocytes is PGC-1α-dependent. Mitochondrial function-associated genes were measured in cultured cardiomyocytes treated with H_2_O_2_ in the absence or presence of hKLF14 overexpression by qPCR. #*P* < 0.05 between the control and H_2_O_2_ groups. ^*^*P* < 0.05 between H_2_O_2_ and the Ad-KLF14+H_2_O_2_ group.

